# Redox control and substrate specificity of the glycogen catabolic isoenzymes in *Synechocystis* sp. PCC 6803

**DOI:** 10.1101/2022.11.21.517384

**Authors:** Niels Neumann, Sofía Doello, Kenric Lee, Bill Kauderer, Karl Forchhammer

## Abstract

Glycogen serves as the main carbon storage polymer in many organisms and is widespread across all domains of life. In cyanobacteria, glycogen degradation is crucial for metabolic transitions during dark phases or during resuscitation from nitrogen starvation. Like many other cyanobacteria, *Synechocystis* sp. PCC 6803 possesses multiple homologues of glycogen catabolizing enzymes, though their specific roles and regulatory mechanisms are only partially understood. Here we demonstrate through biochemical analysis that the glycogen phosphorylase GlgP1, known to promote high temperature acclimation, is uniquely regulated by a C-terminal redox switch found certain cyanobacteria. This is the first evidence of redox regulation of a prokaryotic glycogen degrading enzyme. Notably, GlgP1 is activated via oxidation by reactive oxygen species and inactivated by reducing agents, with thioredoxin being the most effective inhibitor tested. The physiological implications of this redox regulation are discussed. Additionally, a biochemically characterization of the two glycogen debranching isoenzymes GlgX1 and GlgX2 revealed that only GlgX1 exhibits debranching activity, while GlgX2 does not. Mutant analysis confirmed that GlgX1 plays an essential role in glycogen mobilization, being crucial for resuscitation from chlorosis and survival during extended dark periods. In contrast the physiological function of GlgX2 remains unclear.

## Introduction

Glycogen is the primary carbon storage compound in animals, fungi and most bacteria. It is a glucose polymer linked at position α-1,4 with branches at position α-1,6. In bacteria, particularly free-living strains, glycogen is usually built up under conditions of carbon surplus and degraded when energy or carbon availability becomes limited [1–3]. In cyanobacteria glycogen plays a key role in carbon metabolism and is tightly coupled to a photoautotrophic lifestyle: During daylight excess fixed CO_2_ is stored as glycogen which is subsequently broken down at night when metabolism switches to a heterotrophic mode. This essential role is further underscored by the fact that glycogen metabolic enzymes are conserved in all known cyanobacterial strains [4]. Beyond providing energy during dark phases, glycogen degradation is also crucial for acclimation to environmental stress, particularly nitrogen starvation, which is a frequent challenge in marine and terrestrial ecosystems [5]. Adaptation to nitrogen starvation in non-diazotrophic cyanobacteria triggers a genetically determined program, called chlorosis, where cells enter a state of dormancy, enabling them to outlive long periods of nitrogen deprivation. Once combined nitrogen sources become available again, cells can resuscitate within 48 hours, a process primarily fueled by the rapid breakdown of accumulated glycogen which provides the building blocks for re-establishing metabolic processes. [6–8]. Glycogen breakdown requires the coordinated action of glycogen phosphorylase (GlgP) and glycogen debranching enzyme (GlgX): GlgP catalyzes the transfer of orthophosphate to the non-reducing ends of glycogen, releasing glucose-1-phosphate (Glc-1P) which is considered the rate limiting step in glycogen turnover [9]. This processive reaction halts once GlgP reaches the last four glucose residues left on a branch. In prokaryotes those remaining glucose units are hydrolyzed by “direct” debranching enzymes, releasing maltotetraose, which is further processed by maltodextrin phosphorylase (MalP) and by a-1,4-glucanotransferase (MalQ) [10]. The released Glc-1P is converted to Glc-6P by phosphoglucomutase (PGM) enabling its entry into different metabolic pathways including the Embden–Meyerhof–Parnas (EMP), the Entner-Doudoroff (ED) and the oxidative-pentose-phosphate (OPP) pathway.

Like many other cyanobacteria, the unicellular model organism *Synechocystis* sp. PCC 6803 (from now *Synechocystis*) encodes multiple isoenzymes involved in glycogen catabolism, including the glycogen phosphorylases *sll1356* (*glgP1*) and *slr1367* (*glgP2*) as well as the glycogen debranching enzymes *slr0237* (*glgX1*) and *slr1857* (*glgX2*). Previous studies have shown that GlgP2 is responsible for glycogen degradation during the dark and during resuscitation from chlorosis [8]. In contrast, GlgP1 does not contribute to glycogen degradation under those conditions but has been implicated in high temperature acclimation [11]. The roles of GlgX1 and GlgX2 have so far not been investigated and it is unclear whether these enzymes exhibit distinct enzymatic activities or physiological roles in *Synechocystis*. The need for cyanobacteria to switch between autotrophic and heterotrophic metabolic states implies a tight regulation of glycogen catabolism to prevent a premature degradation. While our previous work identified PGM as a key regulator of glycogen metabolism during chlorosis and resuscitation, the divergent roles of GlgP1 and GlgP2 imply additional regulatory mechanisms governing these enzymes [12].

In this study, we address these critical gaps by characterizing the biochemical properties of these isoenzymes, uncovering a novel redox-regulatory switch in GlgP1 and establishing a clear functional hierarchy in glycogen catabolism.

## Results

### Activity of GlgP1 is redox regulated

*Synechocystis* encodes two isoforms of glycogen phosphorylases, the product of *glgP1* (*sll1356*) and *glgP2* (*slr1367*). Previous studies showed that GlgP2 is essential for glycogen degradation during the night and for recovery from nitrogen starvation, whereas GlgP1 appears dispensable under these conditions but was shown to contribute to growth at elevated temperatures [8, 11]. However, the biochemical activity and regulation of GlgP1 and GlgP2 have not been characterized to date.

To determine the enzymatic activity of GlgP1 and GlgP2 we purified the respective recombinant proteins overexpressed in *Escherichia coli.* Since glycogen phosphorylases require pyridoxal phosphate (PLP) as an essential co-factor, we measured UV/Vis absorbance to ensure that both enzymes show equal loading with PLP **(Figure S1)**. GlgP activity was assessed by coupling the formation of glucose-1-phosphate (Glc-1P) from glycogen to the NADPH-producing reactions of phosphoglucomutase (PGM) and glucose-6-phosphate dehydrogenase (G6PDH). Reactions contained 10 mM KH₂PO₄ (saturating concentration; data not shown) and varying amounts of oyster-derived glycogen (0.1–1%) which was used to start the reaction. Both enzymes exhibited robust in vitro activity and were capable of glycogen degradation. While GlgP1 and GlgP2 showed comparable *Kₘ* values for glycogen, GlgP1 displayed a higher overall catalytic efficiency (**Figure 1A and B**, **Table 1**).

**Figure 1:**
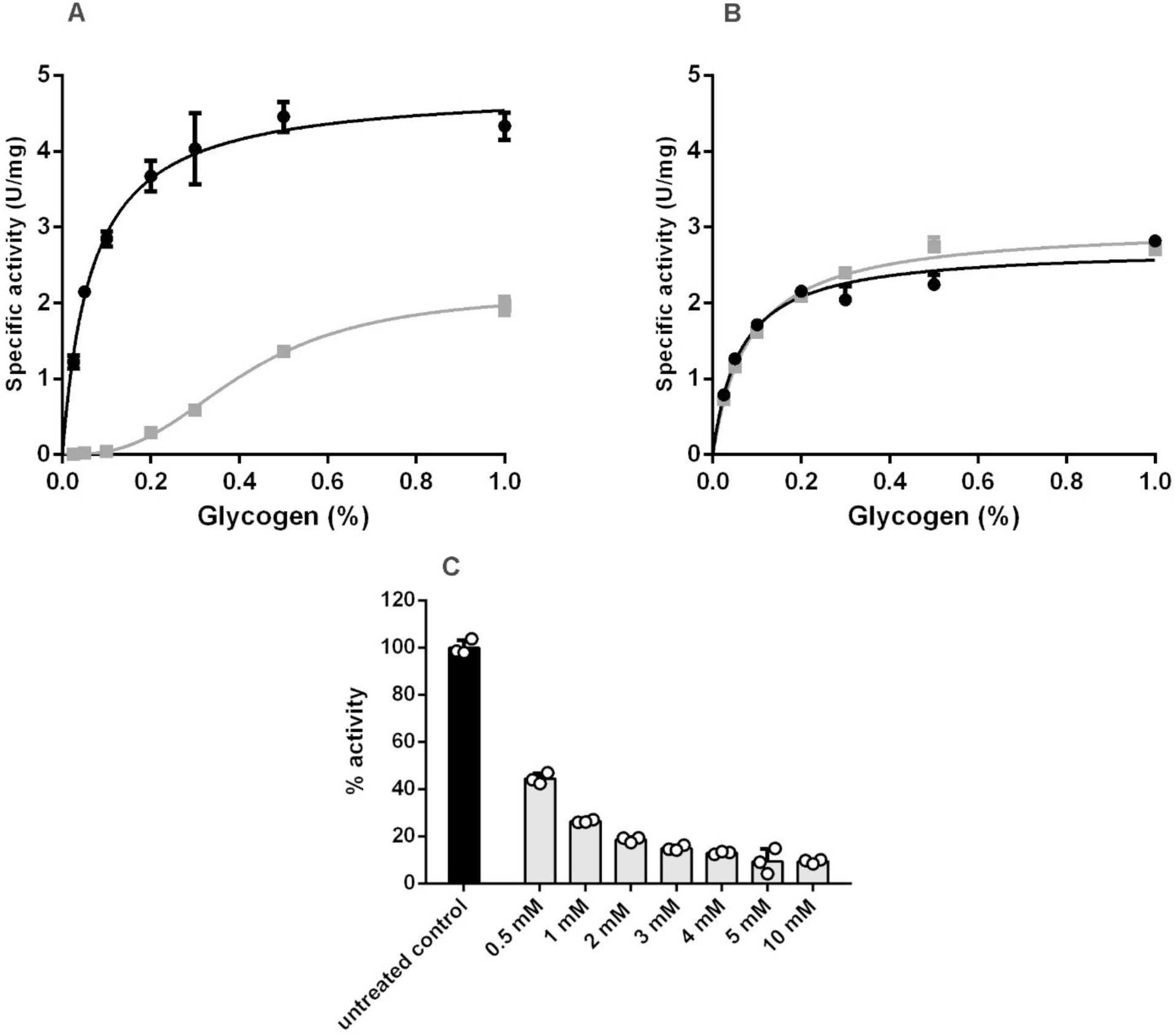
GlgP1 is strongly inhibited under reducing conditions. (A, B) Enzyme kinetics of GlgP1 (A) and GlgP2 (B) with or without 30 min pre-incubation with 5 mM DTT: untreated (black) and DTT-treated (light grey). (C) Relative GlgP1 activity at 0.1% glycogen following 30 min pre-incubation with increasing DTT concentrations, normalized to untreated control (black, set to 100%). Data represent mean ± SD from at least three independent replicates.

**Table 1:** Kinetic parameters of GlgP1 and GlgP2 with and without treatment with 5 mM DTT.

| GlgP1 |  |  |  | GlgP2 |  |  |
| --- | --- | --- | --- | --- | --- | --- |
| Treatment | $k_{cat}(s^{-1})$ | $K_M$ or $K_{0.5}$ (mM) | $K_{cat}/K_M$ or $K_{0.5}$ ($mM^{-1} s^{-1}$) | $k_{cat} (s^{-1})$ | $K_M$ (mM) | $K_{cat}/K_M$ ($mM^{-1} s^{-1}$) |
| Untreated | $6.56 \pm 0.16$ | $0.07 \pm 0.01$ | 93.71 | $4.44 \pm 0.14$ | $0.06 \pm 0.01$ | 74.00 |
| 5 mM DTT | $2.90 \pm 0.08$ | $0.41 \pm 0.09$ | 7.07 | $4.94 \pm 0.09$ | $0.08 \pm 0.01$ | 61.75 |

Values represent the mean ± standard deviation (SD) from triplicate measurements

Inspired by the findings of Florencio, Pérez-Pérez [13] who identified GlgP2 as a putative target of thioredoxin A (TrxA), we investigated whether reducing conditions affect the activity of GlgP1 and GlgP2. Therefore, we pre-incubated enzyme reactions with 5 mM dithiothreitol (DTT) for 30 minutes prior to initiation by glycogen addition. Strikingly under reducing conditions GlgP1 did not follow standard Michaelis-Menten kinetics anymore but showed an allosteric/sigmoidal kinetic curve. Its Catalytic efficiency of GlgP1 dropped to 7.5% of the original activity due to a markedly increased *K_0._*_5_ and a reduced *k*_cat_. In contrast GlgP2 activity remained unaffected by DTT treatment (**Figure 1A and B**, **Table 1**).

To further quantify the redox sensitivity of GlgP1, we performed a DTT titration, incubating the enzyme with 0.5 to 10 mM DTT for 30 minutes. As shown in **Figure 1C**, treatment with 5 mM DTT was sufficient to achieve maximum inhibition of GlgP1 and higher DTT concentrations did not lead to further activity loss. GlgP1 treated with 5 mM DTT was therefore used as a reference for a fully reduced enzyme in all subsequent experiments. Conversely, treatment of the purified enzyme with hydrogen peroxide did not enhance GlgP1 activity, suggesting that the untreated protein is already fully oxidized under native purification conditions (data not shown).

### GlgP1 is selectively inhibited by TrxA and reactivated under oxidizing conditions

To further investigate the redox regulation of GlgP1, we compared the response of GlgP1 and GlgP2 to various reducing agents, including both thiol-dependent and thiol-independent compounds. Enzyme reactions (25 nM GlgP1 or GlgP2) were pre-incubated for 30 minutes with 5 mM DTT, 5 mM tris(2-carboxyethyl)phosphine (TCEP), 5 mM glutathione (GSH), or 5 mM cysteine. In addition, we tested recombinant thioredoxin A (TrxA; product of *slr0623*), applied in its reduced (TrxA_red) or oxidized (TrxA_ox) form at 2.5 µM. Enzymatic activity was assayed using 10 mM KH₂PO₄ and 0.1% oyster glycogen, and untreated enzyme served as a reference control.

Consistent with our previous observations, DTT strongly inhibited GlgP1 activity. Remarkably, TrxA_red also caused substantial inhibition, reducing GlgP1 activity to ∼20% of the untreated control. TCEP, despite having a comparable redox potential (–0.29 V) to DTT (–0.33 V), showed moderate inhibition (∼40% remaining activity). In contrast, GSH, cysteine, and TrxA_ox had no significant effect on GlgP1 **(Figure 2A**). None of the tested treatments, including DTT and TrxA_red, substantially affected GlgP2 activity (**Figure 2B**), underscoring a specific redox sensitivity unique to GlgP1.

**Figure 2:**
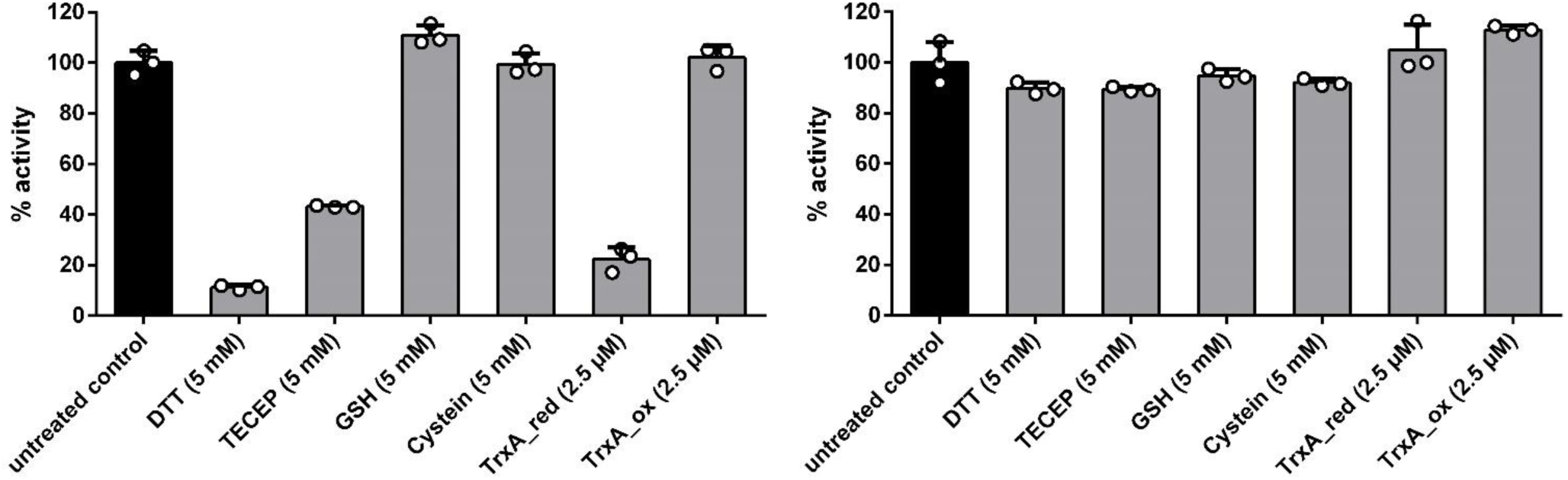
GlgP1 activity is specifically modulated by TrxA. (A) GlgP1 activity after 30 min pre-incubation with various reducing agents (light grey), including DTT, TCEP, GSH, cysteine, and reduced TrxA (TrxA_red); untreated control (black) set to 100%. (B) Same treatments applied to GlgP2. Bars represent mean ± SD of at least three independent replicates.

We next asked whether the inhibition of GlgP1 by reduction could be reversed through oxidation. To this end, GlgP1 was first fully reduced with 5 mM DTT, and subsequently treated with various oxidizing agents for 30 minutes at room temperature. Tested oxidants included oxidized DTT (DTT_ox), glutathione disulfide (GSSG) (5 mM each), and TrxA_ox (10 µM), as well as the stronger oxidants CuCl₂ (5 µM) and H₂O₂ (25 µM), for which optimal concentrations were determined in prior titrations (data not shown).

Compared to the untreated enzyme (set to 100% activity), DTT-reduced GlgP1 retained only ∼10% residual activity. Upon oxidation, H₂O₂ and GSSG showed the strongest reactivation effects, restoring activity to ∼65% and ∼50%, respectively. CuCl₂ treatment led to partial recovery (∼30%), while DTT_ox yielded only modest reactivation (∼20%). TrxA_ox did not increase activity beyond that of the reduced control (**Figure 3**).

**Figure 3:**
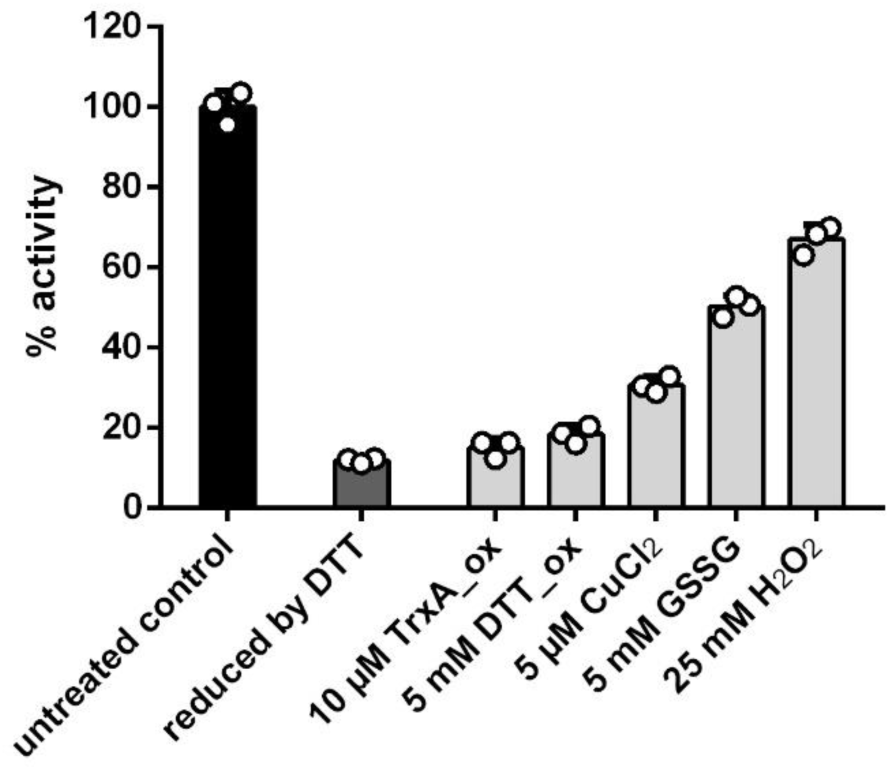
Oxidizing agents restore GlgP1 activity following reductive inhibition. GlgP1 activity under the following conditions: untreated control (black, 100%), DTT-reduced (dark grey), and post-treatment with various oxidants (light grey): oxidized DTT (DTT_ox), GSSG, TrxA_ox, CuCl₂, and H₂O₂. Data represent mean ± SD from at least three replicates.

Together, these results demonstrate that GlgP1 activity is redox-sensitive and can be partially reactivated following reductive inhibition. The pronounced effect of TrxA_red, despite its low concentration compared to the other reducing agents tested, suggests a targeted and specific redox interaction between GlgP1 and TrxA. GlgP2, in contrast, appears largely insensitive to redox modulation under the tested conditions, further highlighting functional divergence between the two isoforms.

### GlgP1 C-Terminal Redox Switch Regulates Enzyme Activity and Diurnal Viability

To identify the regulatory site responsible for redox control of GlgP1 we aligned the amino acid sequence of GlgP1 and GlgP2, revealing a high overall sequence identity of 62%. By looking for prominent cysteine residues which might form disulfide bonds, we discovered a distinctive motif in the C-terminal domain of GlgP1 featuring two cysteine residues at position 837 and 842. Structural prediction using AlphaFold 3 suggests that this region forms a surface exposed hairpin loop potentially stabilized by a disulfide bond between the cysteines C837 and C842 (**Figure 4A**). A comparative sequence analysis across species indicated that this motif is unique to specific cyanobacterial strains (**Figure S2**).

**Figure 4:**
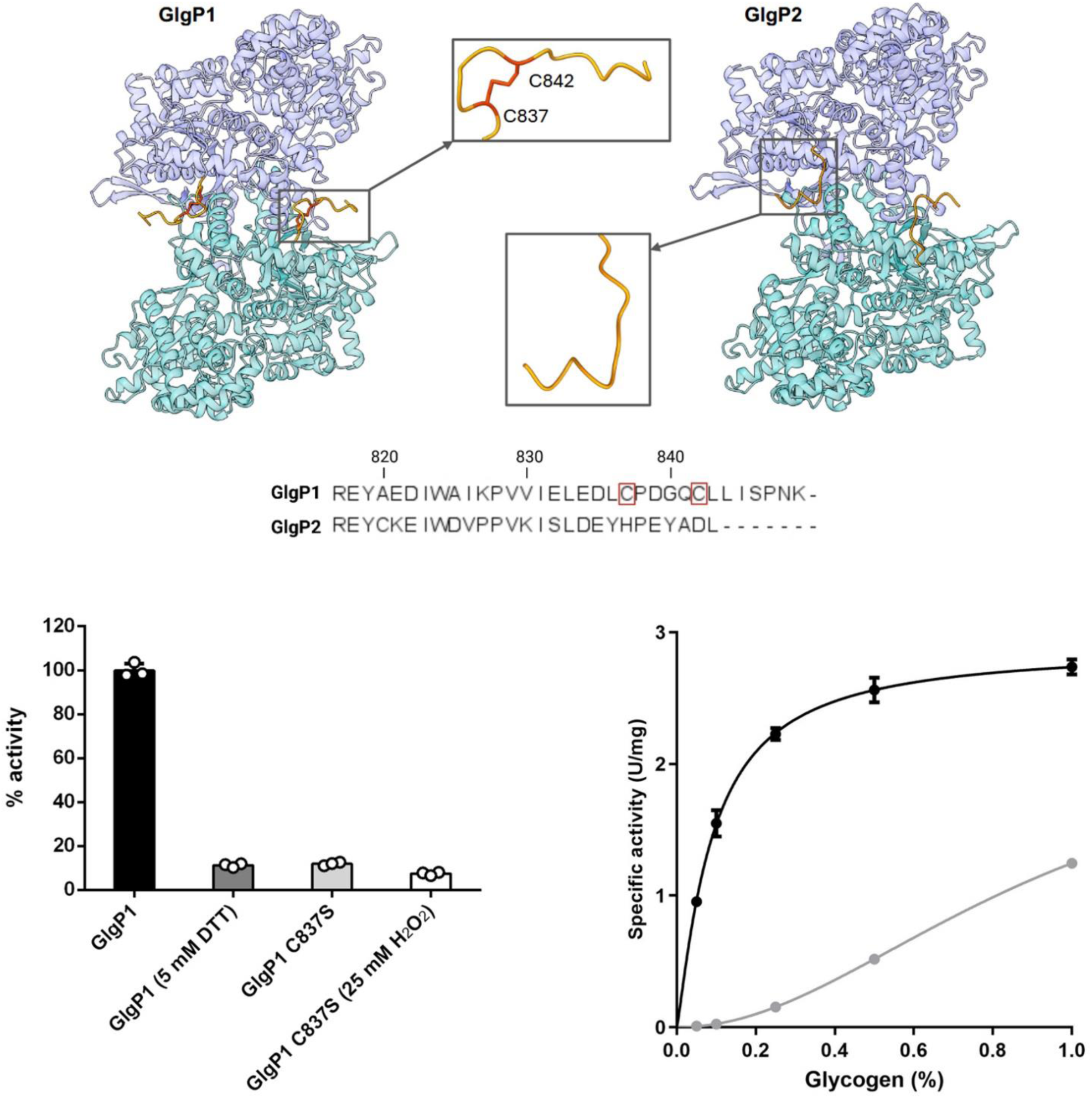
A C-terminal redox switch controls GlgP1 activity. (A) Predicted GlgP1 structure (AlphaFold) highlighting the C-terminal region with Cys837 and Cys842 forming a putative disulfide bond. Sequence alignment shows the absence of these residues in GlgP2. (B) Relative GlgP1 activity: WT (black, untreated = 100%), DTT-reduced WT (dark grey), untreated C837S variant (light grey), and H₂O₂-treated C837S (white). Data represent mean ± SD of at least three replicates. (C) Enzyme kinetics of GlgP1 (black) and GlgP1 (GlgP1ΔC-term)

To assess whether this domain is in fact responsible for redox regulation, we generated a C837S point mutant to mimic a constitutively reduced state. The activity of GlgP1 C837S matched that of the fully reduced GlgP1 WT. However, in contrast to the WT enzyme this variant did not respond to treatment with H_2_O_2_, confirming the redox sensitivity of this cysteine **(Figure 4B)**.

To determine whether the reduced activity of the C837S variant was due to impaired dimerization, we constructed a GlgP1 variant lacking the entire C-terminal domain (GlgP1ΔC-term) and examined its enzymatic activity as well as its oligomeric state via mass photometry. Interestingly, GlgP1ΔC-term displayed enzymatic activity comparable to the C837S mutant, while its ability to dimerize was not negatively affected **Figure 4C, Figure S4A and B**). These findings demonstrate that the C-terminal region of GlgP1 functions as a redox-sensitive regulatory element essential for full enzymatic activity, but not for dimer formation.

To explore the physiological relevance of this redox regulation, we generated *Synechocystis* knockout strains lacking either or both glycogen phosphorylase isoenzymes and assessed their viability under a 12-hour light/dark cycle using a drop-plate assay. While the ***glgP2*** knockout showed a mild growth defect, the ***glgP1*** knockout grew similarly to the wild type (**Figure 5**). However, the growth impairment observed in the ***glgP2*** mutant was significantly worsened in the double knockout, indicating that both isoenzymes contribute to survival under diurnal conditions. These results suggest that GlgP1 can partially compensate for the loss of GlgP2, underscoring the physiological importance of redox regulation of GlgP1 in fluctuating light environments.

**Figure 5:**
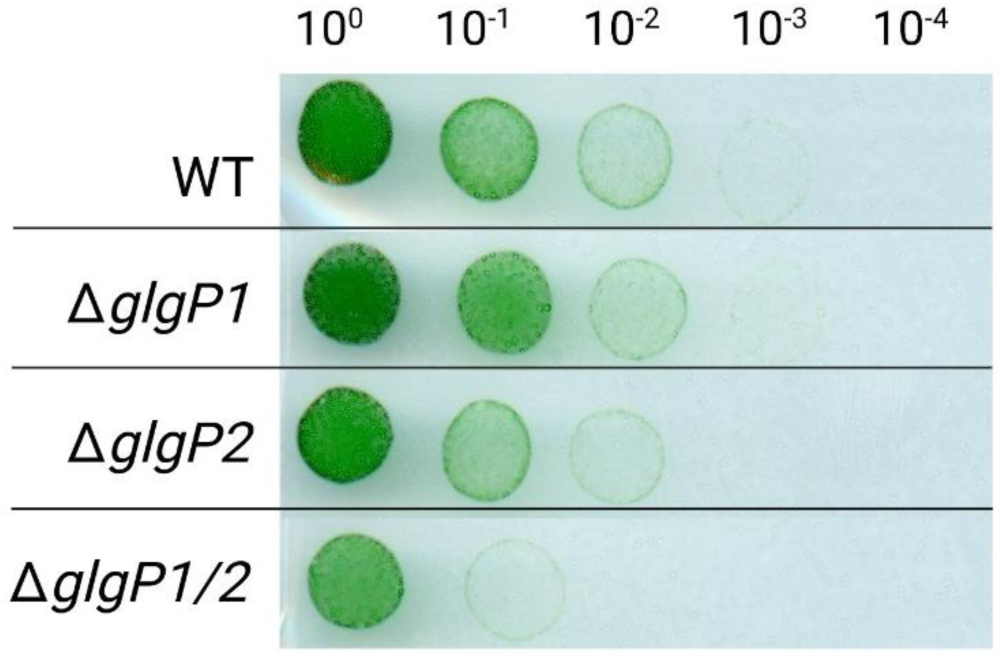
Both glycogen phosphorylases contribute to survival in darkness. Spot assays of *Synechocystis* WT and *glgP* mutants (*ΔglgP1*, *ΔglgP2*, double knockout) under 12 h light/12 h dark cycle. Dilution series indicated in the top row.

### GlgX1 is essential for recovery from chlorosis

Complete glycogen degradation requires not only glycogen phosphorylase activity (GlgP) but also the action of a glycogen debranching enzyme. Similar to GlgP1 and GlgP2, *Synechocystis* encodes two annotated isoforms of glycogen debranching enzymes, *glgX1* (*slr0237*) and *glgX2* (*slr1857*). However, their physiological roles and biochemical activities have not yet been characterized.

To gain insight into their function we generated knockout mutants for both *glgX1* and *glgX2* and assessed their phenotypes under various growth conditions using a drop-plate assay (**Figure 6**). Under continuous light or a 12 h light–dark cycle, both mutants exhibited growth comparable to the wild type (**Figure 6A and B**). However, upon resuscitation from nitrogen chlorosis, the *glgX1* mutant displayed a noticeable growth impairment, whereas the *glgX2* mutant remained unaffected (**Figure 6C**). This phenotype became more pronounced under a 12 h light–dark resuscitation regime (**Figure 6D**) or following extended chlorosis (**Figure 6E**). These results suggest that GlgX1 plays a critical role under conditions that require efficient mobilization of glycogen, specifically during recovery from chlorosis and long-term darkness. In contrast, GlgX1 appears to be dispensable under normal diurnal conditions, where glycogen turnover demands are lower. The physiological role of the paralogue GlgX2 remains elusive, as its deletion did not affect viability under any of the tested conditions.

**Figure 6:**
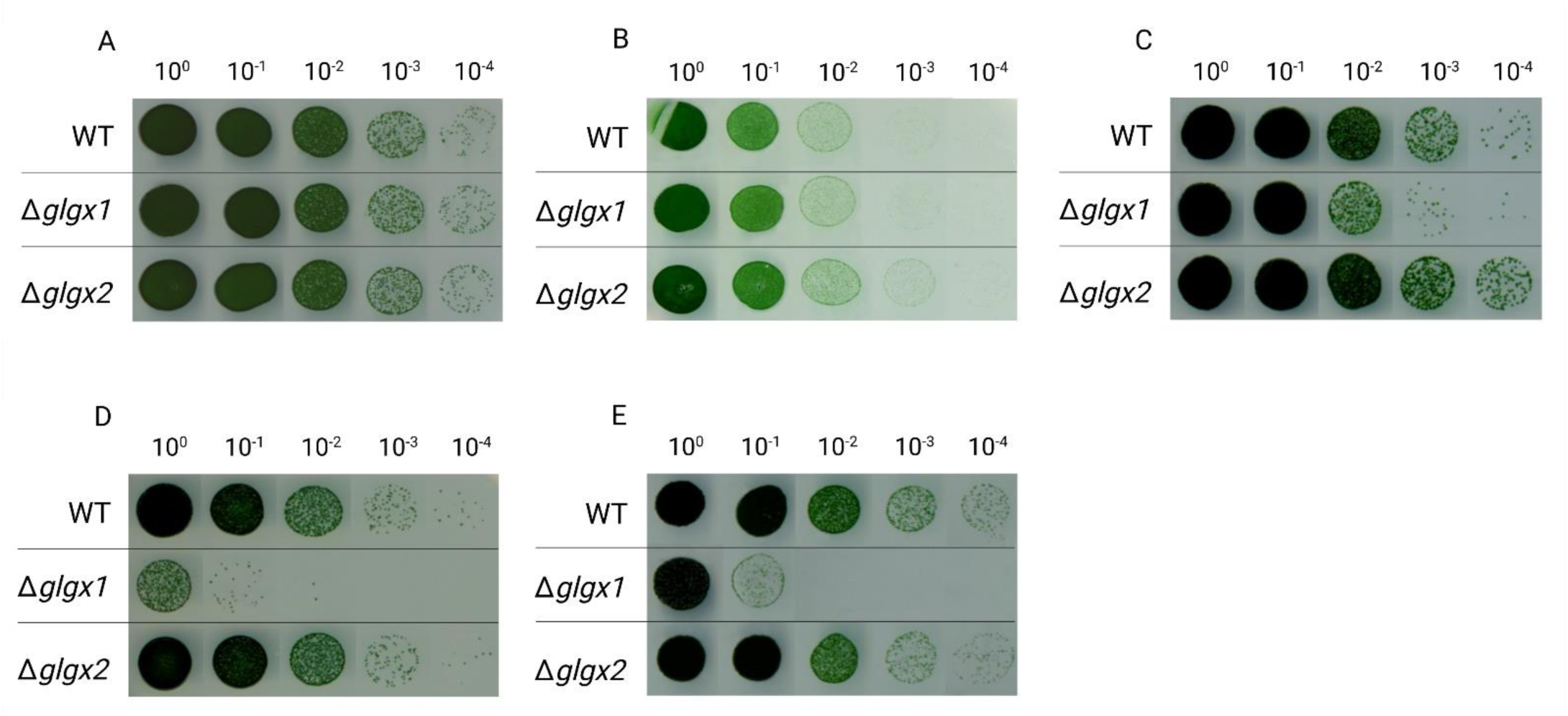
GlgX1 is essential for recovery from chlorosis and survival in darkness. Spot assays of WT, ΔglgX1, and ΔglgX2 strains under various conditions: (A) continuous light, (B) 12 h light/12 h dark cycle, (C) resuscitation after 14 days chlorosis (constant light), (D) resuscitation after 14 days chlorosis (light/dark), (E) resuscitation after 21 days chlorosis (constant light), (F) recovery after 14 days darkness (constant light). Dilution series indicated above.

### GlgX1 promotes GlgP1 and GlgP2 activity, particularly on native glycogen

To further investigate the functional role of GlgX1 and GlgX2, we overexpressed strep-tagged versions of both enzymes in *E. coli* and tested their effects *in vitro* using the established GlgP activity assay. Alongside oyster glycogen, we included glycogen purified from nitrogen-starved *Synechocystis* cultures (both at 0.1% final concentration) to examine how glycogen origin influences enzymatic activity. To facilitate comparison, the activity of GlgP1 and GlgP2 with oyster glycogen and without GlgX addition were set to 100%. Addition of GlgX1 increased GlgP1 activity to 128% with oyster glycogen. When *Synechocystis* glycogen was used as substrate, GlgP1 activity dropped to 65% without GlgX1 but was restored to 117% in its presence (**Figure 7A**). For GlgP2, GlgX1 increased activity to 132% with oyster glycogen and from 40% to 89% with *Synechocystis* glycogen (**Figure 7B**). In contrast, GlgX2 had no significant effect on GlgP activity under any of the tested conditions (**Figure 7A and B**). This experiment yielded two major findings: First, without GlgX1 addition using *Synechocystis* derived glycogen gives rise to generally lower GlgP activities as compared to oyster glycogen (60 % activity for GlgP1 and 40 % activity for GlgP2). Second, addition of GlgX1 enhanced the overall degradation of *Synechocystis* derived glycogen much stronger than degradation of oyster glycogen. While GlgX1 increased activity on oyster glycogen by 28% (GlgP1) and 32% (GlgP2), it boosted activity on *Synechocystis* derived glycogen by 80% and 121%, respectively. Altogether, these results suggest that *Synechocystis* glycogen is more dependent on debranching activity for efficient degradation, and that GlgP2, in particular, benefits strongly from the presence of GlgX1. In contrast, GlgX2 appears to be inactive or non-functional under these assay conditions, further supporting a dominant role for GlgX1 in glycogen catabolism.

**Figure 7:**
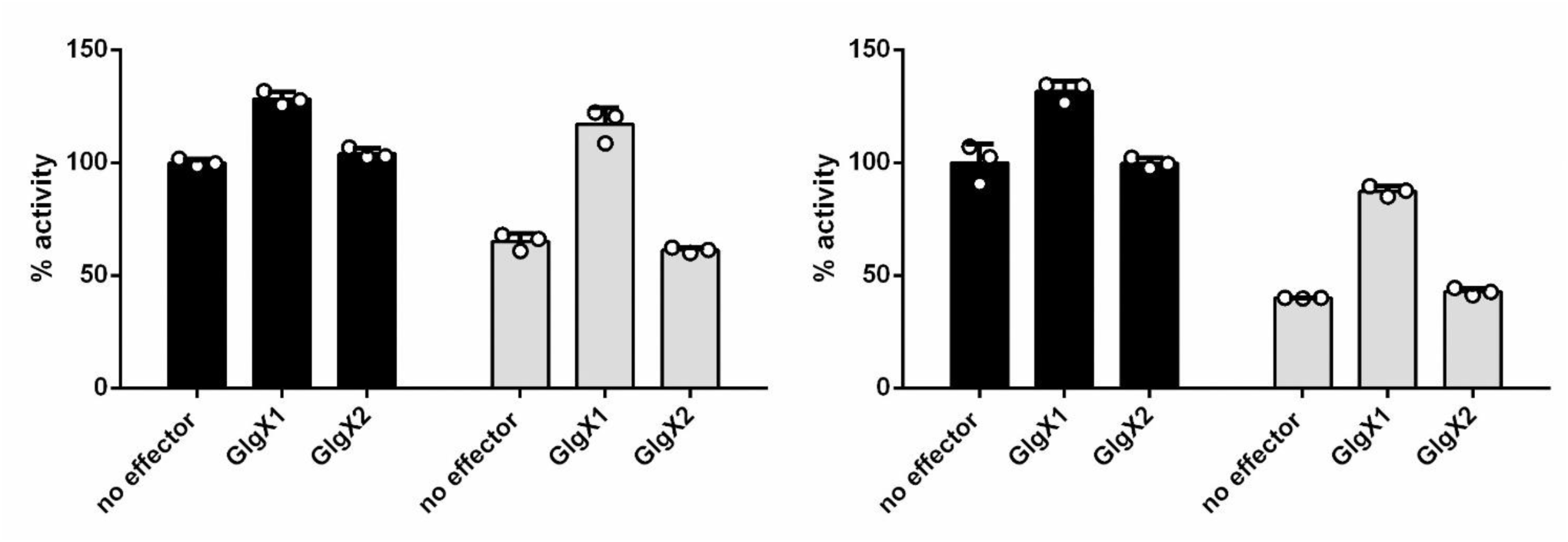
GlgX1 enhances GlgP activity, particularly on Synechocystis-derived glycogen. (A, B) Relative activity of GlgP1 (A) and GlgP2 (B) with or without GlgX1 or GlgX2, using either oyster glycogen (black bars) or Synechocystis glycogen (grey bars). Activities normalized to GlgP alone with oyster glycogen (set to 100%). Data are mean ± SD from at least three replicates.

In addition to testing GlgX activity, we also examined whether GlgX1 or GlgX2 might be subject to redox regulation, as previously shown for GlgP1. To this end, both debranching enzymes were pre-treated with either DTT (5 mM) or H₂O₂ (25 mM) and then added to the GlgP2 assay, using GlgP2 as a redox-insensitive background. Neither GlgX1 nor GlgX2 showed any change in activity following treatment, suggesting that, unlike GlgP1, the debranching enzymes are not redox-regulated under the tested conditions (**Figure S5**).

## Discussion

Glycogen metabolism and its tight regulation is pivotal for cyanobacteria to survive in an ever-changing environment. Understanding the functions and regulatory mechanism of glycogen-associated enzymes is therefore essential for deciphering cyanobacterial physiology. In this study we investigated the glycogen phosphorylase isoenzymes GlgP1 and GlgP2 as well as the glycogen debranching isoenzymes GlgX1 and GlgX2 in *Synechocystis*. We demonstrate that GlgP1 is regulated via a C-terminal redox switch marking the first report of such a mechanism of glycogen phosphorylases in prokaryotes. Additionally, we show that GlgX1 is essential for recovery from chlorosis and enhances glycogen degradation by GlgP activity in dependence of the used glycogen source, whereas the function of GlgX2 remains elusive.

### Redox regulation of GlgP1: A Novel Mechanism in Bacterial Glycogen Phosphorylases

Previous studies established that GlgP2 but not GlgP1 is the primary glycogen phosphorylase responsible for glycogen degradation during the night and resuscitation from nitrogen starvation [8], whereas GlgP1 was shown to play a role in high temperature adaptation[8, 11]. These distinct functions suggested different regulatory mechanisms for the two isoenzymes. Here we confirm that both purified GlgP isoenzymes are active *in vitro,* exhibiting standard Michaelis-Menten kinetics, with GlgP1 displaying a higher overall catalytic efficiency. However, GlgP1 activity is strongly decreased in the presence of reducing agents, adopting a sigmoidal kinetic profile. We identified a unique C-terminal extension in cyanobacterial GlgPs, with GlgP1 containing a conserved cysteine pair (C837 and C842) that forms a regulatory disulfide bond absent in GlgP2. Substitution of C837 with serine resulted in an enzyme activity resembling that of fully reduced GlgP1, underscoring the functional importance of this disulfide bond for enzymatic activation. Truncation of the entire C-terminal domain yielded a similar effect, producing an enzyme with reduced activity comparable to the fully reduced wild-type GlgP1. This suggests that the C-terminal region, and specifically the redox-sensitive cysteines, plays a critical role in regulating GlgP1 activity. The sigmoidal kinetics in the reduced state suggest a diminished glycogen affinity, while an impact on the dimer formation of GlgP1 could be excluded. Although the C-terminal extension appears to be conserved among cyanobacterial GlgPs (Figure S2), the redox-active cysteine motif is restricted to a subset of both diazotrophic and non-diazotrophic strains that are not closely related phylogenetically. This patchy distribution suggests that the motif may have been acquired through horizontal gene transfer rather than by vertical inheritance along a shared evolutionary trajectory. Redox regulation of glycogen metabolizing enzymes has been rarely described in bacteria. A previous study reported redox-dependent modulation of an starch specific amylase in the cyanobacterium *Nostoc* sp. PCC7119 [14]. In humans, an isoenzyme-specific redox regulation of the brain glycogen phosphorylase (bGP) was shown, where the oxidized state affects binding of the activator AMP – an interaction not known to occur in bacterial GlgPs [15]. Unlike bGP, *Synechocystis* GlgP1 is activated under oxidizing conditions, revealing a distinct and previously unrecognized redox-regulatory mechanism in bacterial glycogen metabolism.

### Proposed role of redox -activated GlgP1

The fact that GlgP1 is active only under oxidizing conditions raises questions on the potential role of GlgP1 and under which situations it gets activated. By testing different reducing agents that are present in the cells we could show that reduction and inactivation of GlgP1 was especially effective with TrxA, an m-type thioredoxin which has been shown to be essential in *Synechocystis* [16]. Thioredoxin has been assumed before to play a role in regulation of glycogen metabolism: A proteomic approach by Florencio, Pérez-Pérez [13] revealed that TrxA potentially interacts with several glycogen-associated enzymes in *Synechocystis*, including the glycogen synthatse GlgA2, phorphoglucomutase 1 (PGM1), and the glycogen brancing enzyme (GlgB). Curiously, GlgP1 was not identified in this study. Moreover, the OpcA protein, which regulates the activity glucose-6-phosphate dehydrogenase (G6PDH), is an additional TrxA target. Here, we show that GlgP1 gets re-activated by oxidation, most effectively by H_2_O_2_ and GSSG. While H_2_O_2_ is a common ROS species, GSSG is formed as a product of ROS scavenging by GSH and therefore closely connected to ROS formation. Thereby, a low GSH/GSSG ratio is an indicator for high oxidative stress. Being exclusive to cyanobacteria and the fact that regulation was especially effective with TrxA and H_2_O_2_/GSSG strongly indicates a connection of GlgP1 to photosynthetic activity. TrxA as a part of the ferredoxin/thioredoxin system is known to provide the biochemical link between light reactions and the regulation of carbon metabolism in photoautotrophic organisms [17]. Reactive oxygen species are byproducts of the photosynthetic electron transport chain, whose formation increases with light intensity and temperature, a feature encountered by all photosynthetically active organisms.

We also showed that both glycogen phosphorylases are able to contribute to night-time survival, with *glgp2* being the isoenzyme that likely contributes more to glycogen degradation in the dark, consistent with its expression levels. Despite being downregulated during the night, residual *glgp1* could also contribute to glycogen degradation in the dark, and might also be metabolically important for the transition from night to day. A previous study addressed the potential role of GlgP1, showing that GlgP1 is important for survival at high temperatures and that GlgP1 is only active during illumination [11]. High temperature has been shown to cause strong ROS formation in bacteria and in nature high temperature and high light are often associated with one another [18]. High temperatures have a negative influence on photosynthesis by increasing the oxygenase activity of RuBisCo and due to the fact that CO_2_ shows a lower water solubility at higher temperatures than O_2_ [19]. This increases the demand of ribulose-1,5-BP due to a higher rate of photorespiration. Several studies have shown that glycogen degradation can contribute to replenish the ribulose-1,5-BP pool by shuffling carbon through the OPP pathway to stabilize photosynthesis [20]. This mechanism appears to be especially important during the “restart” of photosynthesis after the night phase, where it has been shown that glycogen degradation at the end of the dark phase can fill up the depleted ribulose-1,5-BP pools, leading to a faster start of photosynthesis while glycogen free mutants show a strong delay [21]. We assume that the same pathway contributes to ribulose-1,5-BP formation during heat stress. Furthermore, it was shown that survival at higher temperature and high ROS formation is dependent on the availability of GSH [22, 23]. Reduction of GSSG by glutathione reductase is dependent on NADPH, which can either be formed by linear photosynthetic electron transport or in the OPP fueled by glycogen catabolism. The oxygen-evolving complex of PSII has also been shown to be prone to heat damage, which would inhibit NADPH formation by linear electron transport, making the OPP the only source of NADPH [24].

### GlgX1 is the key glycogen debranching enzyme in *Synechocystis*

It is estimated that roughly one-third of a glycogen granule is accessible to glycogen phosphorylase (GlgP) without the need to remove α-1,6-linked branches [25]. Therefore, glycogen debranching enzymes are crucial for a complete glycogen degradation. Despite their importance, little is known about their function in cyanobacteria, and the roles of the two GlgX isoenzymes in *Synechocystis* have remained unexplored. Here, we identify GlgX1 (Slr0237) as the primary glycogen debranching enzyme in *Synechocystis*. While neither the *glgX1* nor *glgX2* knockout strains showed growth defects under standard diurnal conditions, the *glgX1* mutant was severely impaired in surviving prolonged darkness. This suggests that during normal night periods, partial glycogen degradation by glycogen phosphorylases is sufficient to meet basal metabolic needs. In contrast, when glycogen reserves must be mobilized for extended periods, as during prolonged darkness, debranching activity becomes essential for accessing the full glycogen pool **(Figure 6**). A pronounced phenotype was also observed for the *glgX1* mutant during recovery from nitrogen starvation, particularly after extended chlorosis or under day-night resuscitation. This highlights the distinct metabolic demands of resuscitation, which relies not just on energy generation but also on anabolic processes requiring carbon skeletons. Glycogen degradation during resuscitation thus appears more dependent on complete access to stored glycogen via GlgX1 activity, whereas nocturnal metabolism may tolerate limited breakdown.

In contrast, *glgX2* deletion had no detectable effect under any tested condition, and GlgX2 showed no activity *in vitro*. Debranching enzymes are notoriously difficult to annotate with confidence, as their glycoside hydrolase domains can overlap functionally with those of transglycosylases or other carbohydrate-active enzymes. Thus, GlgX2 may possess a yet-undiscovered specificity or function unrelated to classical debranching activity.

### Cooperativity of GlgP and GlgX depends on enzyme isoform and glycogen architercture

Our *in vitro* assays confirmed that GlgX1 enhances GlgP activity, but this effect strongly depended on the source of glycogen. Enhancement was more pronounced with glycogen isolated from *Synechocystis* than with commercially available oyster glycogen, suggesting that the structure of native glycogen is more reliant on debranching activity for efficient degradation. Notably, GlgP2 activity showed a greater dependence on GlgX1 than GlgP1, but only when *Synechocystis* derived glycogen was used. This could reflect differences in branching pattern or other structural features specific to *Synechocystis* glycogen. In eukaryotes, glycogen is known to undergo post-synthetic modifications that influence degradability and a smilar complexity might exist in cyanobacteria [26]. *Synechocystis* possesses two glycogen synthases, both contributing to glycogen biosynthesis [27]. Previous studies have shown that glycogen branching patterns differ slightly between single synthase mutants, whereas the wild type contains a mixture of both [28]. Future work should explore whether GlgP1 and GlgP2 show different preferences or efficiencies depending on which synthase produced the glycogen.

In contrast to GlgP1, GlgX1 and GlgX2 did not exhibit any detectable redox regulation (**Figure S5**), indicating that the primary level of control during glycogen breakdown lies with the GlgP enzymes themselves. The preferential stimulation of GlgP2 by GlgX1, combined with the similar resuscitation phenotypes observed in the respective knockout mutants, suggests that GlgP2 and GlgX1 function as a coordinated enzymatic pair during metabolic recovery from chlorosis. Further investigation is needed to uncover potential regulatory mechanisms governing this GlgP2-GlgX1 axis. Nonetheless, our findings provide new insight into the functional specialization of glycogen catabolic enzymes in cyanobacteria and highlight the sophisticated regulation that underpins carbon reserve utilization in *Synechocystis*. Whether these regulatory principles extend to other cyanobacterial species or are conserved across broader phylogenetic boundaries remains an open question for future research.

## Methods

### Cultivation of *Synechocystis*

All *Synechocystis* sp. PCC 6803 strains used in this study were grown in BG_11_ supplemented with 5 mM NaHCO_3_, as described previously [29]. A list of the strains used in this study is provided in **Table S1**. Cultivation was performed with continuous illumination (40–50 μmol photons m^−2^ s^−1^) and shaking (130–140 rpm) at 27°C if not stated otherwise. Induction of nitrogen starvation and resuscitation was induced as described previously [30, 31]. If mutants or strains containing antibiotic markers were used, the precultures were propagated with the appropriate concentration of antibiotics. Biological replicates were inoculated with the same precultures but propagated, nitrogen starved, and resuscitated independently in different flasks under identical conditions.

### Isothermal, Single-Reaction DNA Assembly

Cloning was performed as described by Gibson, Young [32] using E. coli NEB10β cells (details in Table S2). All primers and plasmids used are shown in **Table S3** and **Table S4** in the supplemental material, respectively.

### Cultivation of Escherichia coli

If not otherwise stated *E. coli* was grown in Luria-Bertani medium at 37°C. For growth on plates, 1.5% (w/v) agar-agar was added. For cells containing plasmids, the appropriate concentration of antibiotics was used. All *E. coli* strains used in this study are listed in **Table S2**.

### Protein purification

The plasmids used for protein overexpression are shown in **Table S4**. *Escherichia coli* Rosetta-gami (DE3) (details **Table S2**) was used for the overexpression of all proteins. All purified proteins were tagged at the C-terminus with a Strep-tag. For overexpression, cells were cultivated in 2xYT (3.5% tryptone, 2% yeast extract, 0.5% NaCl; 1L of culture in 5L flasks) at 37 °C until reaching exponential growth (OD_600_ 0.6-0.8). Protein overexpression was induced by adding 75 µg/L anhydrotetracycline, followed by incubation at 20°C for 16 h. Cells were harvested by centrifugation at 4000 g for 10 min at 4 °C. Cell disruption was performed by sonication in 40 mL of lysis buffer (100 mM Tris-HCl pH 8, 150 mM NaCl, 10 mM MgCl_2_). Lysis buffers were supplemented with DNAse I, and cOmplete^™^ protease inhibitor cocktail (Roche, Basel). The cell lysate was centrifuged at 40,000 g for 45 min at 4°C and the supernatant was filtered with a 0.22 µM filter.

For purification of the strep tagged proteins a 5 mL Strep-Tactin XT column (GE Healthcare, Illinois, USA) was used. The cell extracts were loaded onto the column with a peristaltic pump followed by washing with washing buffer (50 mM Tris-HCl pH 8.0, 150 mM NaCl) and elution with elution buffer (50 mM Tris-HCl pH 8.0, 150 mM NaCl, and 50 mM biotin).

### GlgP and GlgX enzymatic assays

Buffer for enzymatic reactions was composed of 50 mM HEPES-KOH pH 7.5, 10 mM MgCl_2_, 1 mM NADP^+^, 5 mM KH_2_PO_4_, 1U/µl phosphoglucomutase from rabbit muscle (P3397, Merck), 10 µM Glucose-1,6-bisphosphate and 1 U/mL G6PDH from *Saccharomyces cerevisiae* (G6378, Merck). For GlgP1/2 activity, 2,5 µg of Strep-tagged purified protein was added to each reaction. Reaction was started by the addition of glycogen. Reactions were carried out in a total of 300 µl in a 96-well microplate. Absorption change at 340 nm was continuously measured for 15 min at 30 °C in a TECAN Spark® Multiplate reader (Tecan Group AG, Männedorf, Switzerland). For determining GlgX activity, 15 µg of GlgX1 and GlgX2 Strep-tagged purified protein were added to the GlgP assay mastermix respectively.

### Redox Treatments

For reduction, purified enzymes were incubated with the indicated concentrations of reducing agents for 30 minutes at room temperature, followed by enzymatic activity measurement as described above.

For reoxidation experiments, GlgP1 was first reduced with 5 mM DTT for 30 minutes to ensure full reduction of cysteine residues. Excess DTT was subsequently removed using a HiTrap Desalting Column (Cytiva) connected to a peristaltic pump and pre-equilibrated with buffer (50 mM Tris-HCl, pH 7.5, 150 mM NaCl). Eluted protein fractions were immediately used for oxidation. For reoxidation, the reduced GlgP1 protein was incubated for 30 minutes at room temperature with one of the following oxidizing conditions: 25 mM hydrogen peroxide (final ∼30 mM), 10 µM oxidized thioredoxin A (TrxA), 5 mM oxidized DTT, 5 µM CuCl₂, or 5 mM oxidized glutathione (GSSG). Residual oxidants were removed by a second buffer exchange using the same desalting column setup. Enzymatic activity was subsequently assayed under standard conditions.

### Mass Photometry Analysis

Protein samples were diluted with buffer (50 mM Tris-HCl pH 7.5, 5 mM MgCl2) to aconcentration of 100 nM. Microscope coverslips (No. 1.5H, 24×5, Marienfeld, Germany) were cleaned 3 times by immersion in isopropanol (HPLC grade) and Milli-Q water. CultureWell gaskets (3 mm DIA x 1 mm Depth, Grace Bio-Labs, Bend, OR, USA) were assembled on the clean coverslips. 5 μL of buffer were added to a gasket’s well, the focal position was identified and secured in place with an autofocus system based on total internal reflection for the entire measurement. 5 μL of diluted protein were then added to the same well and movies of 60 s duration were recorded. Each sample was measured at least three times independently. A calibration measurement was performed to correlate contrast with mass. Images were processed and analyzed using the DiscoverMP v2022 R1 software as described in [33]

### Spot viability assay

Serial dilutions of chlorotic cultures were prepared (10^0^, 10^−1^, 10^−2^, 10^−3^, 10^−4^, and 10^−5^), starting with an OD_750_ of 1.5 µL of these dilutions, dropped on solid BG_11_ agar plates, and cultivated at 50 μmol photons m^−2^ s^−1^ and 27°C for 5 to 7 d.

### Statistical analysis

Statistical details for each experiment can be found in the figure legends. GraphPad PRISM was used to perform one-sided ANOVA to determine the statistical significance. Asterisks (*) in the figures symbolize the p-value: One asterisk represents p ≤ 0.05, two asterisks p ≤ 0.01, three asterisks p ≤ 0.001, and four asterisks p ≤ 0.0001

## Data availability statement

All data are included in this manuscript or in supportive information

## Author CRediT Statement

**Niels Neumann,** investigation, methodology, formal analysis, visualization, writing original draft. **Sofia Doello: conceptualization, supervision, review and editing. Kenric Lee**: investigation. **Bill Kauderer**: investigation. **Karl Forchhammer,** conceptualization, validation, supervision, writing – review and editing, project administration, funding acquisition.

## Acknowledgments

This work was supported by the DFG funded research consortium FOR2816 “The Autotrophy-Heterotrophy Switch in Cyanobacteria: Coherent Decision-Making at Multiple Regulatory Layers”. We also acknowledge infrastructural funding via the Cluster of Excellence (EXC2124) “Controlling Microbes to Fight Infections”

## Supporting Figures

**Figure S1:**
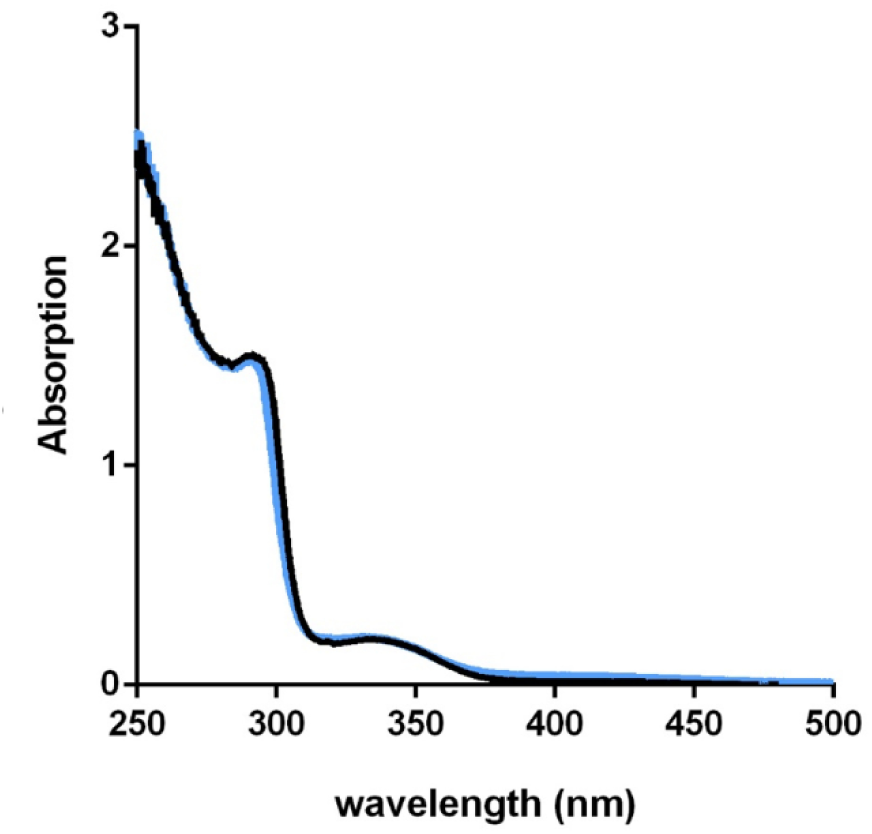
PLP loading confirmed for GlgP1 and GlgP2 by absorbance spectra. UV/Vis spectra of purified GlgP1 (black) and GlgP2 (blue). Absorbance at 280 nm reflects aromatic residues; absorbance at 335 nm indicates PLP in its enolimine form.

**Figure S2:**
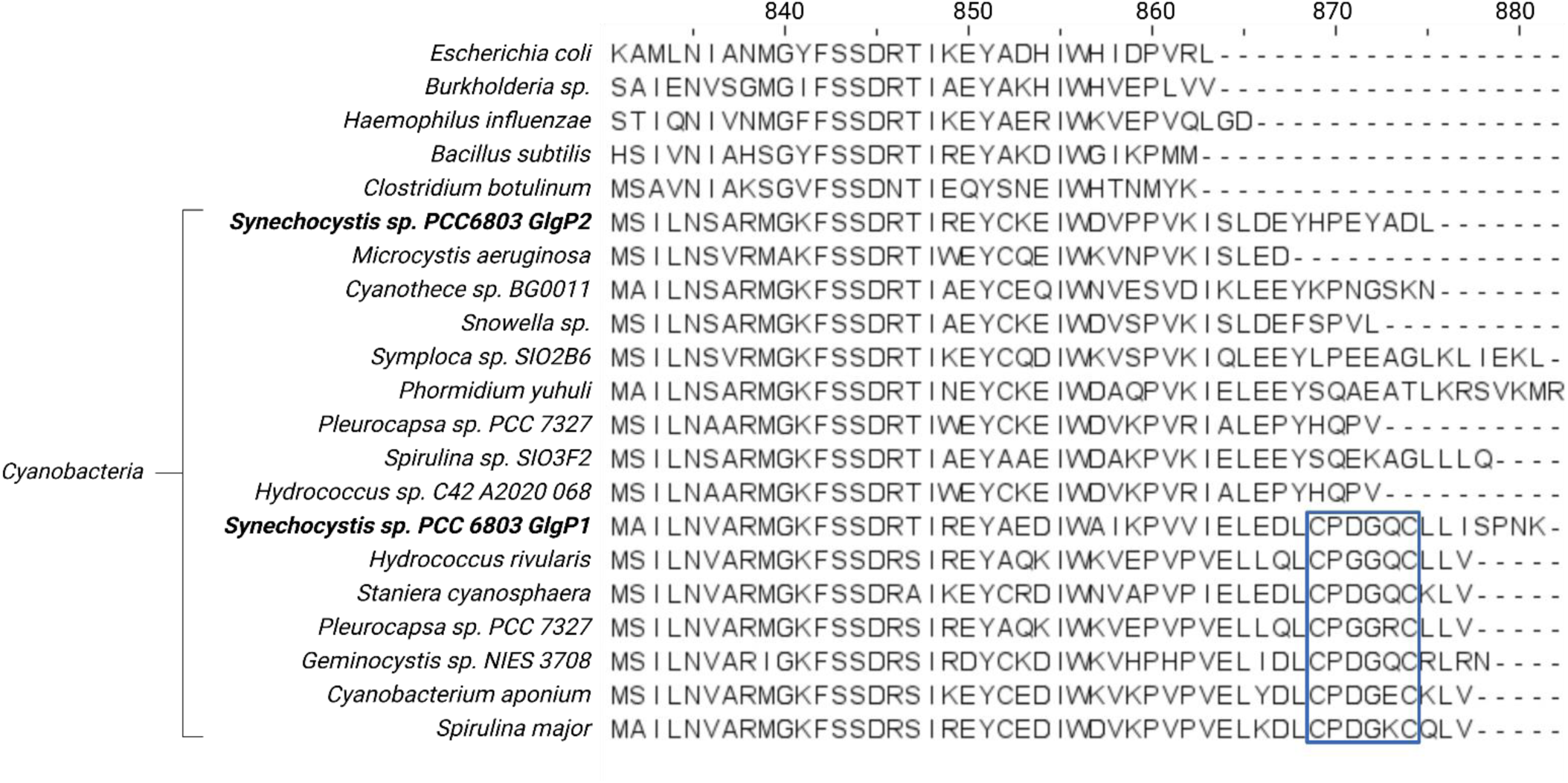
The GlgP1 redox switch is conserved in select cyanobacteria. Sequence alignment of the GlgP1 C-terminal region from various bacterial species. Conserved cysteine motif in GlgP1 homologues is highlighted a blue box

**Figure S3:**
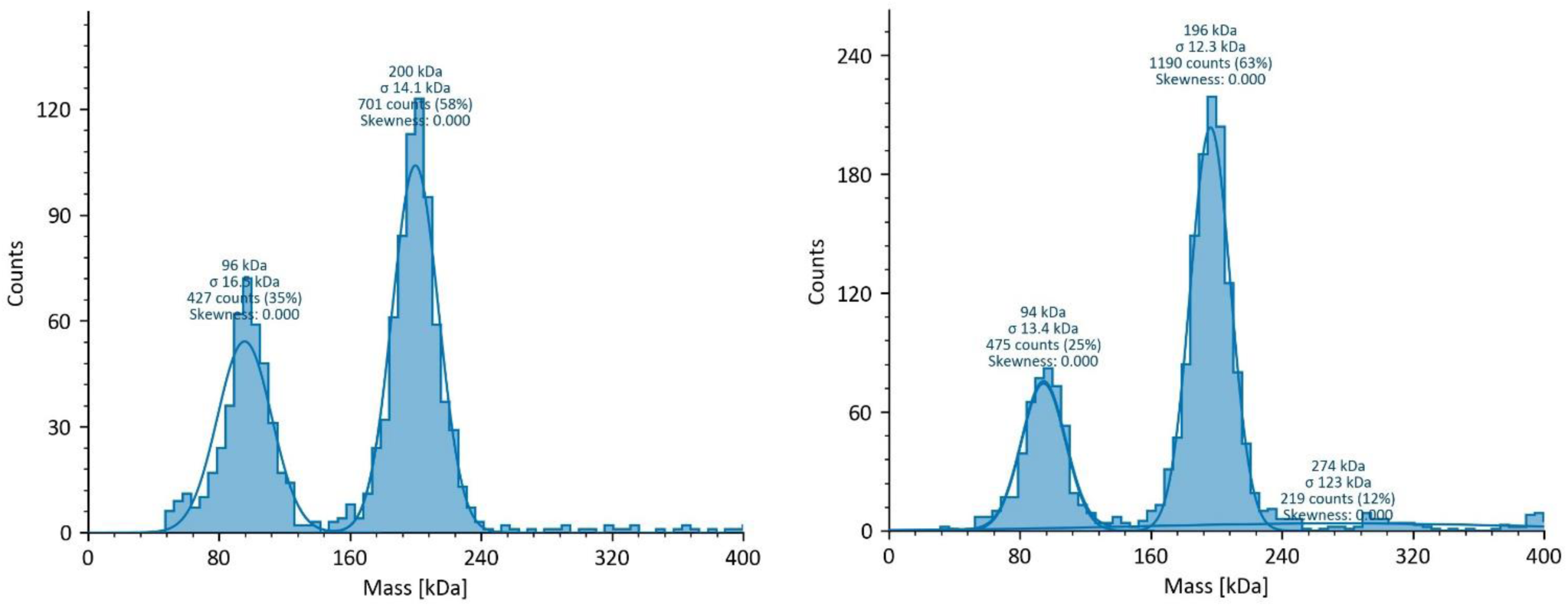
Mass photometry analysis of the oligomeric state of WT GlgP1 ang GlgP1ΔC-term.

**Figure S4:**
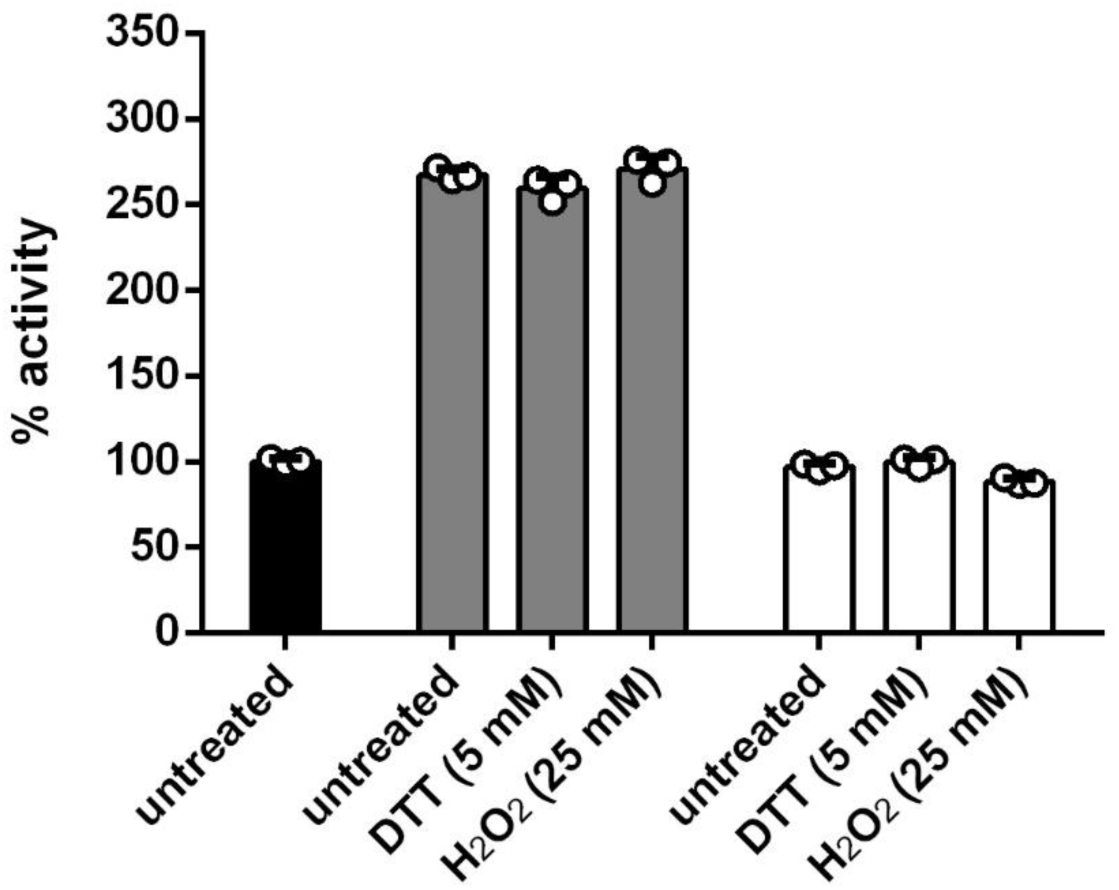
GlgX1 and GlgX2 activity is not influenced by redox state. Relative GlgP2 activity in the presence of GlgX1 or GlgX2 pre-treated with either DTT or H₂O₂. Control reactions without GlgX (dark grey), with GlgX1 (light grey), and with GlgX2 (white). *Synechocystis* glycogen was used as substrate. Data are mean ± SD of at least three replicates

## Supporting tables

**Table S1:**
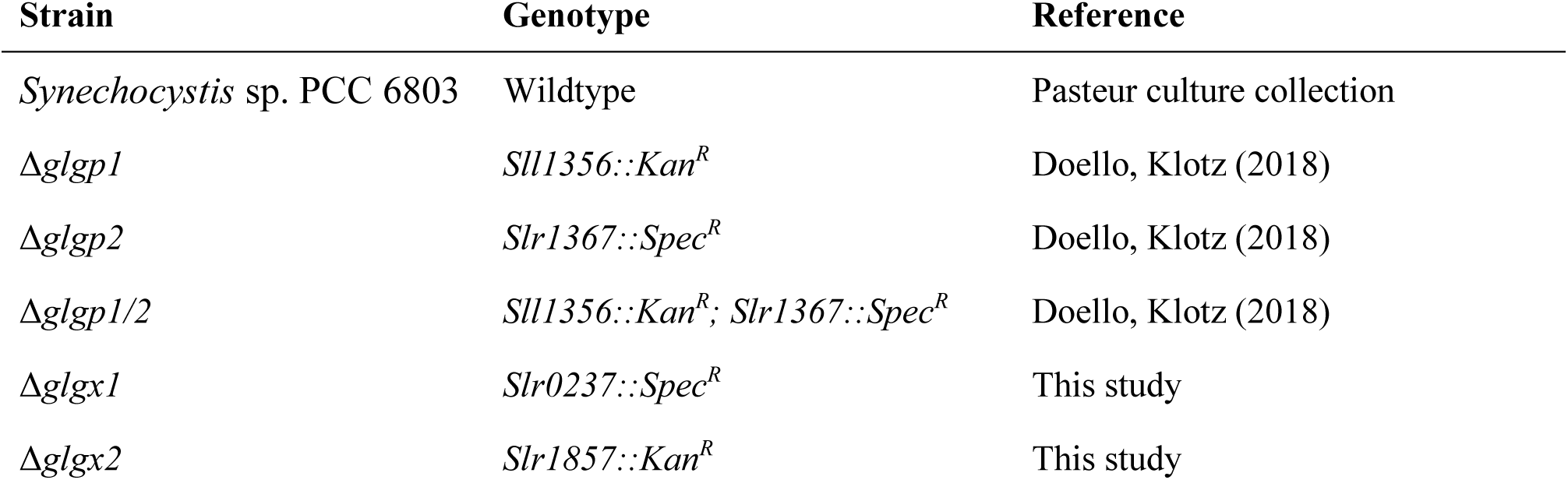
List of cyanobacterial strains used in this study.

**Table S2:** List of strains used for cloning and protein overexpression.

| Strain | Genotype | Description |
| --- | --- | --- |
| *E. coli* NEB10β | Δ(ara-leu) 7697 araD139 fhuAΔlacX74 galK16 galE15 e14-Φ80dlacZΔM15 recA1 relA1 endA1 nupGrpsL(StrR) rphspoT1 Δ(mrr-hsdRMS-mcrBC) | Molecular cloning |
| *E. coli* Rosetta gami (DE3) | Δ(ara-leu)7697 ΔlacX74 ΔphoA PvuII phoR araD139 ahpC galE galK rpsL (DE3) F'[lac<sup>+</sup> lacI<sup>q</sup> pro] gor522::Tn10 trxB pLysSRARE (Cam<sup>R</sup>, Str<sup>R</sup>, Tet<sup>R</sup>) | Protein overexpression |

**Table S3:**
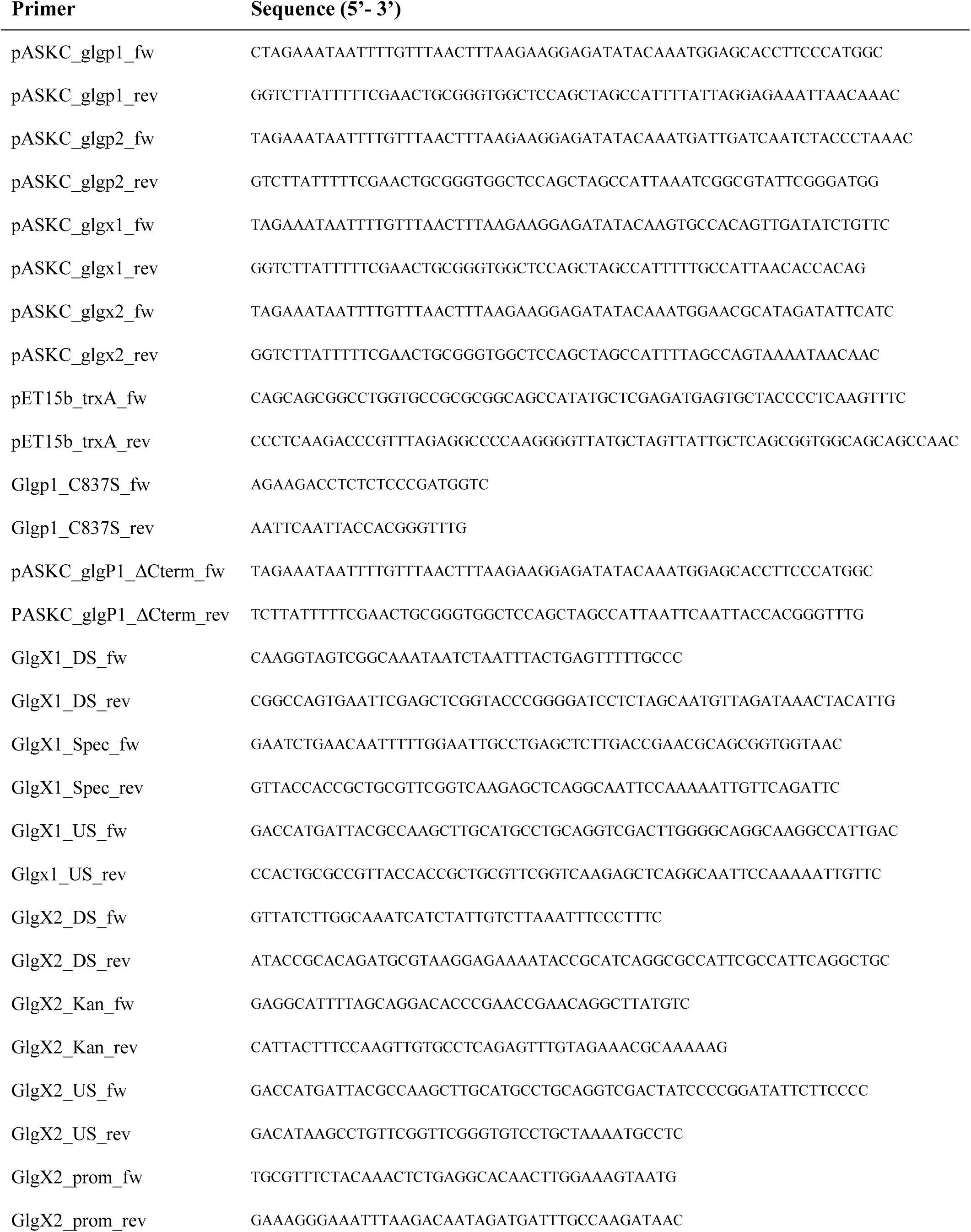
List of primers.

**Table S4:** List of used plasmids.

| Plasmid Name | Description | Resistance Marker | Source/Reference |
| --- | --- | --- | --- |
| pASK IBA5plus | Template for pASKC construction | Ampicillin | IBA Lifesciences (Göttingen, Germany) |
| pASKC | Overexpression with C-terminal Strep-tag | Ampicillin | This study |
| pASKC_glgp1 | Expression of C-terminal Strep-tagged *Synechocystis* GlgP1 in *E.coli* | Ampicillin | This study |
| pASKC_glgp2 | Expression of C-terminal Strep-tagged *Synechocystis* GlgP2 in *E.coli* | Ampicillin | This study |
| pASKC_glgp1_C837S | Expression of C-terminal Strep-tagged *Synechocystis* GlgP1 C837S in *E.coli* | Ampicillin | This study |
| pASKC_glgp1ΔC-term | Expression of C-terminal Strep-tagged *Synechocystis* GlgP1ΔC-term in *E.coli* | Ampicillin | This study |
| pASKC_glgx1 | Expression of C-terminal Strep-tagged *Synechocystis* GlgX1 in *E.coli* | Ampicillin | This study |
| pASKC_glgx2 | Expression of C-terminal Strep-tagged *Synechocystis* GlgX2 in *E.coli* | Ampicillin | This study |
| pASKC_trxA | Expression of C-terminal Strep-tagged *Synechocystis* TrxA in *E.coli* | Ampicillin | This study |
| pUC19_glgx1_KO | Replacement of *slr0237* for Spec<sup>R</sup> in *Synechocystis* sp. PCC 6803 | Ampicillin | This study |
| pUC19_glgx2_KO | Replacement of *slr1857* for Kan<sup>R</sup> in *Synechocystis* sp. PCC 6803 | Ampicillin | This study |

